# Multi-ancestry genome-wide analysis identifies common genetic variants for self-reported sleep duration and shared genetic effects

**DOI:** 10.1101/2022.02.16.480716

**Authors:** Ben H Scammell, Yanwei Song, Cynthia Tchio, Takeshi Nishiyama, Tin L Louie, Hassan S Dashti, Masahiro Nakatochi, Phyllis C Zee, Iyas Daghlas, Yukihide Momozawa, Jianwen Cai, Hanna M. Ollila, Susan Redline, Kenji Wakai, Tamar Sofer, Sadao Suzuki, Jacqueline M Lane, Richa Saxena

**Affiliations:** Center for Genomic Medicine, Massachusetts General Hospital and Harvard Medical School, Boston MA, USA; Program in Medical and Population Genetics, Broad Institute, Cambridge MA, USA; Department of Anesthesia, Critical Care and Pain Medicine, Massachusetts General Hospital, Boston MA, USA; Cardiovascular Research Institute, Morehouse School of Medicine, Atlanta GA, USA; Department of Public Health, Nagoya City University Graduate School of Medicine, Nagoya, Japan; Department of Biostatistics, University of Washington, Seattle, WA, USA; Public Health Informatics Unit, Department of Integrated Health Sciences, Nagoya University Graduate School of Medicine, Nagoya, Japan; Center for Circadian and Sleep Medicine, Department of Neurology, Feinberg School of Medicine, Northwestern University, Chicago, IL, USA; Laboratory for Genotyping Development, RIKEN Center for Integrative Medical Sciences, Yokohama, Japan; Department of Biostatistics, University of North Carolina at Chapel Hill, NC, USA; Department of Medicine, Harvard Medical School, Boston, MA, USA; Division of Sleep and Circadian Disorders, Brigham and Women’s Hospital, Boston, MA, USA; Department of Preventive Medicine, Nagoya University Graduate School of Medicine, Nagoya, Japan

**Author notes:** These authors jointly supervised the work Corresponding Author: Richa Saxena, PhD Center for Genomic Medicine, Massachusetts General Hospital, 185 Cambridge Street, CPZN 5.802, Boston, MA, 02114, USA. **Abbreviations**: CHARGE (Cohorts for Heart and Aging Research in Genomic Epidemiology), EUR (European), EAS (East Asian), AHL (Admixed Hispanic/Latino), AFR (African), SAS (South Asian), eQTL (expression quantitative trait loci), UKBB (UK Biobank), HCHS/SOL (Hispanic Community Health Study/Study of Latinos), J-MICC (Japan Multi-Institutional Collaborative Cohort Study).

**Keywords:** self-reported sleep duration, genome-wide association studies, multi- ancestry meta-analysis, polygenic scores, genetic transferability

## Abstract

Both short and long sleep duration are associated with increased risk of chronic diseases, but the genetic determinants of sleep duration are largely unknown outside of European populations. Here we report transferability of a polygenic score of 78 European ancestry sleep duration SNPs to an African (n=7,288; p=0.003), a South Asian (n=7,485; p=0.025), and a Japanese (n=13,618; p=6x10^-4^) cohort, but not to a cohort of Hispanic/Latino (n- XXX; p=0.71) participants. Furthermore, in a pan-ancestry (*N* = 483,235) meta-analysis of genome-wide association studies (GWAS) for habitual sleep duration, 5 novel and 68 known loci are associated with genome-wide significance. For the novel loci, sleep duration signals colocalize with expression-QTLs for PRR12 and COG5 in brain tissues, and pleiotropic associations are observed with cardiovascular and neuropsychiatric traits. Overall, our results suggest that the genetic basis of sleep duration is at least partially shared across diverse ancestry groups.

## Introduction

Both short (≤6 h per night) and long (≥9 h per night) sleep duration are associated with psychiatric illness, cardiovascular disease, type 2 diabetes, obesity, and all-cause mortality (1–3). Twin studies have identified a heritable component of habitual sleep duration (*h^2^* = 0.3-0.6) (4–6), and functional studies have linked rare, missense mutations in *BHLHE41* to habitually short sleep (7). Genome-wide association studies (GWAS) have uncovered many genetic loci associated with self-reported sleep duration (8–11), and in the largest GWAS of sleep duration to date, Dashti and colleagues identified 78 independent associations in a cohort of European ancestry (*n* = 446,118) (10). Findings have been consistent across cohorts; for example the *PAX8*, *VRK2*, and *FBXL12/UBL5/PIN1* loci replicated in the Cohorts for Heart and Aging Research in Genomic Epidemiology (CHARGE) Consortium study (n = 47,180) and a polygenic score of all 78 loci associated with 0.66 (0.54 to 0.78) mins/allele greater sleep duration in the CHARGE study (8). Dashti et al also identified robust genetic correlations between sleep duration and disorders such as schizophrenia (*r*g = 0.26), depression (*r*g = 0.31), and type 2 diabetes (*r*g = 0.28) (10).

In studies of the US population, sleep duration varies by over an hour between groups of divergent genetic ancestry (12–14). Understanding the genetic basis of sleep duration in non-European populations is important for understanding these disparities. For example, the *PAX8* signal was identified in Europeans and replicated in a cohort of African-American individuals (rs1191684, *p* = 9.3 × 10^−4^, n=4,771) (8). In a meta-analysis of Japanese subjects, the *PAX8* and *VRK2* loci were both insignificant, but a novel association at the *ALDH2* locus (rs671 **G/**A, *p* = 1.2 × 10^−8^, *n*=31,230) was identified (11).

The *ALDH2* variant is associated with caffeine (15) and alcohol (16) intake, but may be specific to East Asian groups as rs671 A is very rare (MAF < 0.01) in non-East Asians (17). The genetic basis of sleep duration in non-European populations is poorly understood, mainly due to under-representation of diverse ancestry groups in biobanks (18, 19).

Delineating the genetics of sleep duration in diverse, non-European populations will help to identify novel genetic factors influencing sleep physiology and help to ensure equitable translation of scientific advances from genomics across populations. An important step in this direction is to assess the transferability of known genetic loci (20). To address this issue, we studied the cross-ancestry transferability of polygenic scores (PGS) and individual variants from the EUR population. Then we conducted a meta-analysis of five sleep duration GWAS of diverse ancestries (Admixed Hispanic/Latino (AHL), African (AFR), East Asian (EAS), European (EUR), and South Asian (SAS)) and identified five novel loci. Lastly, we assessed these loci for associations with other sleep and psychiatric phenotypes and colocalized the associations with eQTL signals from 13 brain regions and pituitary tissue assessed by GTEx (21).

## Results

### Transferability of EUR associations to non-European genetic ancestries

To extend GWAS of sleep duration to non-EUR ancestry groups, we first identified sub- groups of South Asian (SAS; *n* = 7,485) and African (AFR; *n* = 7,288) genetic ancestry from the UK Biobank (UKBB) using k-means clustering (k=4) (**Fig. S1**, **Table S1**). We performed GWAS for self-reported sleep duration in each sub-group adjusting for age, sex, 10 principal components (PCs) of ancestry, genotyping array, and genetic relatedness (**Table S2**). GWAS was also conducted in Admixed Hispanic/Latinos (AHL; *n* = 8,726) from HCHS/SOL for average sleep duration calculated from self-reported bed and wake times (see **Methods**), which adjusted for age, sex, 5 PCs of ancestry, study center, relatedness, and Hispanic background group. GWAS results were also obtained from two previous habitual sleep duration association studies (European (EUR) from UKBB, *n* = 446,118 (10) and Japanese (EAS) from J-MICC, *n* = 13,618) (11), and cohort characteristics were tabulated for the five GWAS (**Table 1**). Mean self-reported sleep duration ranged from 6.6 (SD=1.0) h per night in EAS to 7.9 (SD=1.4) h per night in AHL. Narrow-sense heritability (*h*^2^) was calculated for each cohort with GCTA (22) and ranged from 7.3% in AFR to 15.0% in EUR (**Table S2**).

**Table 1:** Study Cohort characteristics.

| **Ancestry** | **European (EUR)** | **East Asian (EAS)** | **Admixed Hispanic/Latino (AHL)** | **African (AFR)** | **South Asian (SAS)** |
| --- | --- | --- | --- | --- | --- |
| Biobank | UK Biobank | J-MICC | HCHS/SOL | UK Biobank | UK Biobank |
| N | 446,118 | 13,618 | 8,726 | 7,288 | 7,485 |
| Age, yrs (SD) | 57.30 (8.00) | 54.60 (9.40) | 46.16 (14.08) | 51.82 (8.06) | 53.39 (8.46) |
| Female, N (%) | 245,073 (54.1) | 7,449 (54.7) | 5,249 (60.2) | 4,125 (57.9) | 2,760 (46.1) |
| BMI, kg/m<sup>2</sup> (SD) | 27.4 (4.7) | 23.1 (3.3) | 29.84 (6.03) | 29.4 (5.4) | 27.3 (4.4) |
| Sleep Duration, h per night (SD)<sup>a</sup> | 7.20 (1.10) | 6.60 (1.00) | 7.94 (1.38) | 6.71 (1.37) | 6.79 (2.33) |
| Shift workers, N (%)<sup>b</sup> | 23,556 (5.2) | N/A | 0 | 826 (11.6) | 599 (10.0) |
| Smoking status: Current, N (%) | 47,112 (10.4) | 2,547 (18.7) | 1618 (18.5) | 891 (12.5) | 725 (12.1) |
| Alcohol intake, drinks/week (SD) | 2.86 (1.50) | 2.0 (2.7) | 5.49 (8.13) | 3.20 (1.88) | 2.27 (1.70) |

The five single-ancestry groups derive from the UK Biobank (UKBB), the Japan Multi-Institutional Collaborative Cohort Study (J-MICC), and the Hispanic Community Health Study/Study of Latinos (HCHS/SOL). Cohort characteristics were calculated following cohort selection and sample quality control exclusions.
<sup>a</sup>UKBB and J-MICC participants were asked to self-report sleep duration, whereas HCHS/SOL participants self-reported habitual bed and wake times on weekdays and weekends from which average nightly sleep duration was calculated.
<sup>b</sup>The shift work measure includes participants who reported “usually” or “sometimes” working shifts. HCHS/SOL participants who reported shift work, extreme sleep duration (<3 or >18 h per night), and sleep medication use (n=4,077) were excluded during the selection of the AHL cohort.

We then assessed the transferability of the EUR associations in the non-European GWAS cohorts. We first constructed a PGS using 78 independent lead variants associated with sleep duration in the EUR study (10) and tested for associations in the four non-EUR ancestry GWAS using GTX (Genetics ToolboX, R package) (**Table 2**) (23). The PGS was significantly associated with sleep duration (*p* < 0.05) in the AFR, EAS, and SAS ancestry cohorts, and effects on nightly sleep duration ranged from 0.04-0.83 minutes per copy of the effect allele (min/allele) (**Table 2**). Notably, the PGS was not significant in the AHL cohort (*p* = 0.71). This may be due to demographic differences between the AHL and EUR GWAS cohorts, how sleep duration was assessed in HCHS/SOL study (calculated from self-reported bed and wake times from weekday and weekend), or genetic differences between European and Hispanic/Latino populations (24).

**Table 2:** Transferability 78 SNP Polygenic Scores from EUR ancestry to non-EUR cohorts.

| Cohort | *N* | *Beta*<br>(min/allele) | 95% CI<br>(min/allele) | *P*<sup>a</sup> | *P* (het)<sup>b</sup> |
| --- | --- | --- | --- | --- | --- |
| AFR | 7,288 | 0.62 | [0.41 - 0.83] | **2.8 × 10<sup>-3</sup>** | 0.100 |
| AHL | 8,726 | 0.07 | [-0.29 - 0.43] | 7.1 × 10<sup>-1</sup> | 0.195 |
| EAS | 13,618 | 0.38 | [0.22 - 0.54] | **6.2 × 10<sup>-4</sup>** | 0.195 |
| SAS | 7,485 | 0.39 | [0.04 - 0.74] | **2.5 × 10<sup>-2</sup>** | 0.683 |

Sleep duration PGS of 78 SNPs from EUR ancestry to non-EUR cohorts. The PGS from the European ancestry GWAS was significant in the African, East Asian, and South Asian GWAS cohorts and predicted changes in habitual sleep duration of 0.04-0.83 minutes per effect allele copy (min/allele). Statistics were calculated with the Genetics ToolboX (GTX) R package.
<sup>a</sup>*P* is the test statistic for the 1 d.f. chi-squared test of the risk score from the base population (EUR) to the target population (non-EUR cohort). Significant PGS tests (*P* < 0.05) are bolded.
<sup>b</sup>*P* (het) is the test statistic for the heterogeneity test, which tests effect size differences between the base (EUR) and target (non-EUR) cohorts.

Second, to understand the relationship between transferability and minor allele frequency (MAF), we plotted the effect sizes and MAF of the 10 most significant signals from the EUR GWAS (*p* < 2.4 × 10^−11^) across all ancestry groups (**Fig. 1**) (19). As rare alleles may be recent and thus population-specific, the transferability of a locus can be better understood by considering the MAF of lead variants (19). We found that the minor alleles of lead variants at *PAX8*, *FTO*, and *PPP2R3A* were common in all cohorts and their effects were highly transferable. The effect sizes of the *PAX8* minor allele (rs6737318 G, MAF > 0.11 across ancestry groups) in AHL, EAS, and SAS (1.74 [0.96] min/allele - 2.22 [1.74] min/allele) were similar to that in EUR subjects (2.46 [0.18] min/allele). In the AFR cohort, rs6737318 was associated with a relatively large effect on sleep duration (7.86 [1.98] min/allele, MAF=0.11) and was significant after correction for multiple testing (*p* = 7.3 × 10^−4^) (**Table S3**). As expected, some variants with low MAF showed limited transferability (20). For example, the minor alleles rs13109404 G (*BANK1*; all non-EUR MAF < 0.03) and rs78797502 T (*VRK2*; AFR MAF = 0.02; AHL MAF = 0.08; SAS MAF = 0.11) were uncommon in the non-EUR cohorts, and their effect sizes were relatively inconsistent across cohorts.

**Figure 1:**
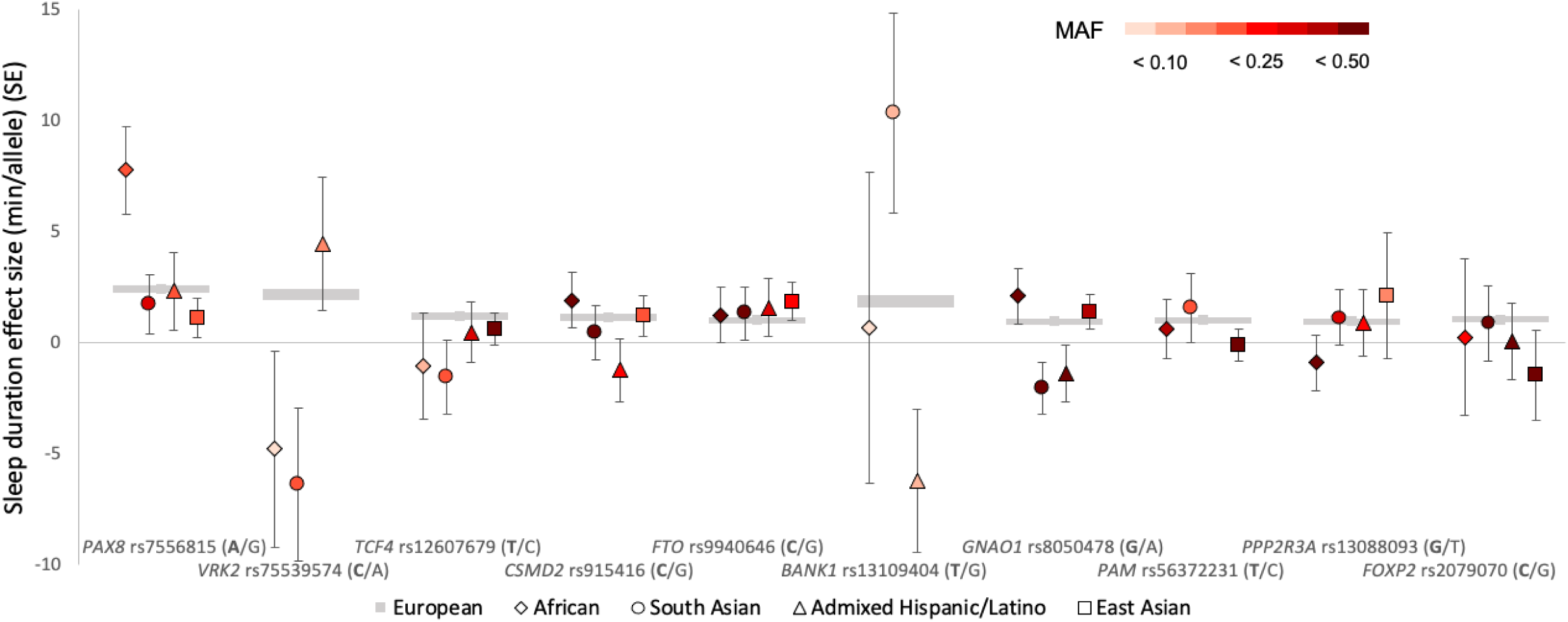
Cross-ancestry transferability of the 10 most significant (*p* < 2.4 × 10^−11^) EUR loci. Cross-ancestry transferability of the 10 most significant (*p* < 2.4 × 10^−11^) EUR loci. The effect sizes for the 10 most significant EUR loci are shown for the EUR GWAS (gray) and four non-EUR GWAS. Minor allele frequencies (MAF) range from 0.0 (absent) to 0.59. All cohorts are sign-concordant for the *PAX8* (rs7556815) and *FTO* (rs9940646) signals; for the other loci at least one non-EUR cohort was not sign- concordant with the EUR signal.

Third, to understand the importance of the aggregate burden of the polygenic EUR signal, we constructed a genome-wide PGS (from EUR GWAS) and assessed transferability to AFR and SAS ancestral groups from the UK Biobank, for which individual-level data was available (**Fig. 2**). Using PRSice (25), we found that transferability was greatest (with maximum model fits) at the p=0.061 PGS threshold for the AFR cohort (*R^2^* = 0.0037) and the p=0.003 PGS threshold for the SAS cohort (*R^2^* = 0.0039) (**Fig. 2a, 2c**). This suggests that many variants which are not genome-wide significant (*p* > 5.0 × 10^−8^) contribute to the genetic architecture of sleep duration. Last, an increasing EUR genome-wide polygenic burden predicted increased sleep duration; we found that higher quartiles of genetic burden predicted greater effects on self-reported duration, and Q4 predicted the greatest effect on sleep duration in AFR (14.9 [4.9] minutes) and SAS (9.4 [4.7] minutes) relative to Q1 (**Fig. 2b, 2d**). These findings suggest that the genetic basis of sleep duration is at least partially conserved across divergent ancestries.

**Figure 2:**
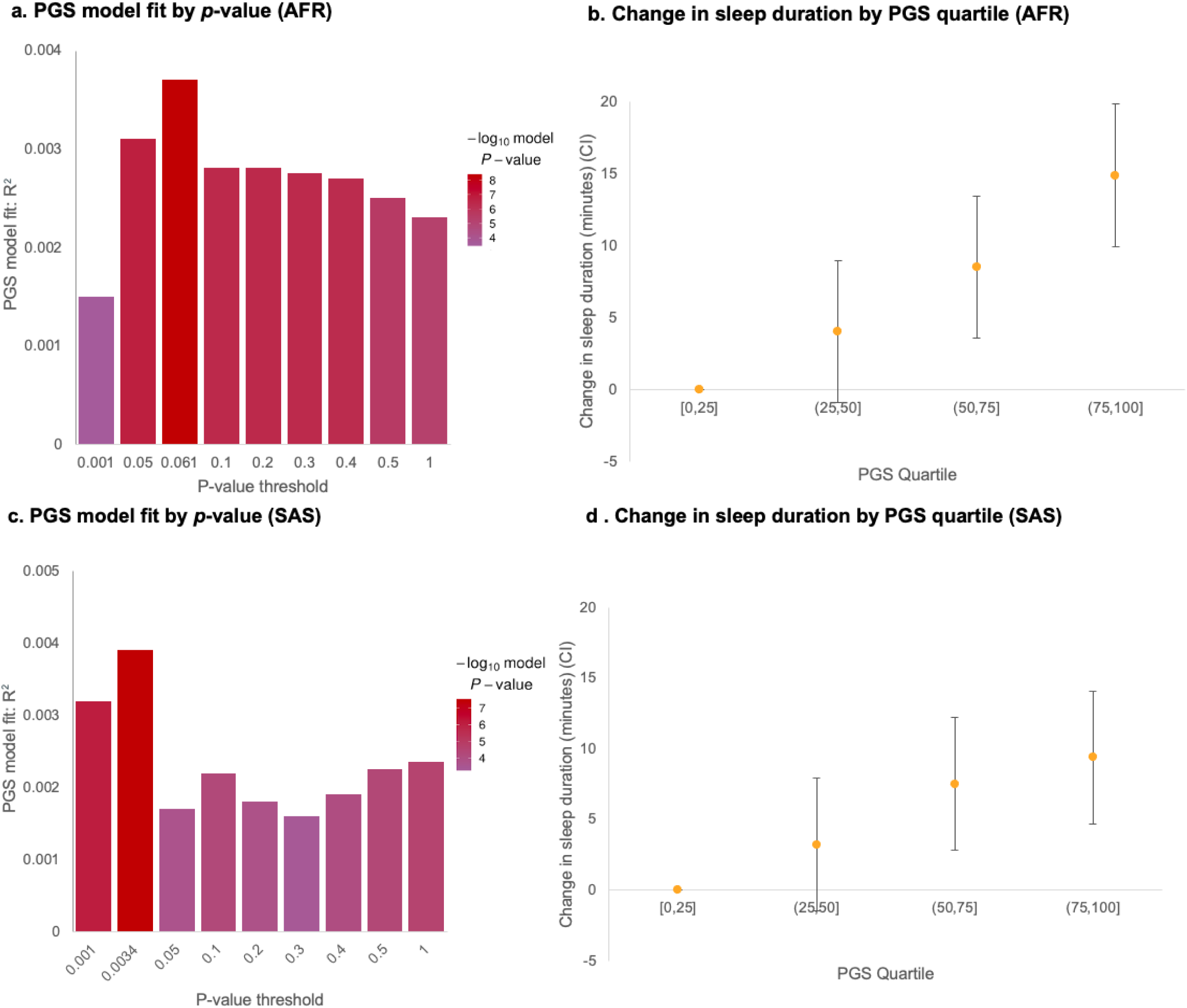
Transferability (*R^2^*) and PGS quartile plots for AFR and SAS cohorts. Transferability (*R^2^*) and PGS quartile plots for AFR and SAS cohorts. **a** & **c**, PGS model fit (*R^2^*) at multiple p-value thresholds for the AFR and SAS cohorts. The maximum observed transferability for AFR was at *p* = 0.061 (*R^2^* = 0.0037) and at *p* = 0.0034 (*R^2^* =0.0039) for the SAS cohort. **b** & **d**, Change in sleep duration by polygenic score quartile for AFR and SAS. The confidence interval associated with the [0,25] strata was designated as a reference factor in PRSice (Methods).

### Multi-ancestry meta-analysis identifies novel sleep duration loci

To identify novel sleep duration associations in a multi-ancestry cohort, we conducted a fixed-effects meta-analysis of the five GWAS (AFR, n=7,288; AHL, n=8,726; EAS, n=13,618; EUR, n=446,118; SAS, n=7,485) using METAL (26). In this analysis, we examined 19,794,469 SNPs from 483,235 individuals (**Fig. 3**), and we identified 73 independent genome-wide significant (*p* < 5.0 × 10^−8^) signals which include five novel genetic associations and 68 associations previously identified in the EUR study (**Table 3; Table S3**) (10). Two of the association signals (rs11928693 near *HACD2* and rs75639044 near *COG5*) had been observed in a GWAS of European ancestry UKBB samples using a subset that excluded related individuals (*n* = 384,371) (27). As expected, significance at the 73 loci was driven primarily by SNP associations from the European ancestry GWAS, but notably, directions of effect for lead SNPs were highly concordant across ancestral groups; effect directions were consistent across at least 4 of the 5 cohorts for 52/73 (71.2%) lead SNPs (binomial p = 1.1 × 10^−4^), and the effect directions for the 5 novel signals were consistent across all cohorts (binomial p = 3.1 × 10^-2^, **Table S3**). The mean effect size of the 73 genome-wide significant signals was 1.00 (SE=0.16) minute per copy of the effect allele. A second fixed-effects meta-analysis for the non-European ancestry cohorts (AFR, AHL, EAS, and SAS cohorts, *n* = 37,117) did not identify additional loci (**Fig. S2**, λ = 1.035).

**Table 3:** Novel genome-wide significant loci discovered in multi-ancestry meta-analysis.

|  |  |  |  |  | European<br>ancestry GWAS<br>(*n* = 446,118) |  | non-EUR meta-analysis<br>(*n* = 37,117) |  | Multi-ancestry<br>meta-analysis<br>(*n* = 483,235) |  |  |
| --- | --- | --- | --- | --- | --- | --- | --- | --- | --- | --- | --- |
| Lead SNP | Chr:Bp<br>(GRCh37) | Nearest<br>gene<sup>a</sup> | Alleles<br>(E/A)<sup>b</sup> | EAF<br>range<sup>c</sup> | Beta<br>(SE)<sup>d</sup> | *P* | Beta<br>(SE) | *P* | Beta<br>(SE) | *P* | Effect<br>Direction |
| rs11928693 | 3:123,275,179 | *HACD2* | T/G | 0.03-0.53 | 0.77<br>(0.18) | $3.80 \times 10^{-7}$ | 1.71<br>(0.76) | $2.48 \times 10^{-2}$ | 0.84<br>(0.18) | $4.49 \times 10^{-8}$ | +++++ |
| rs7689303 | 4:170,203,771 | *SH3RF1* | G/T | 0.22-0.61 | 0.72<br>(0.12) | $3.00 \times 10^{-7}$ | 1.14<br>(0.54) | $1.51 \times 10^{-2}$ | 0.72<br>(0.12) | $3.22 \times 10^{-8}$ | +++++ |
| rs6909225 | 6:73,608,356 | *KCNQ5* | A/G | 0.16-0.34 | 0.72<br>(0.12) | $2.80 \times 10^{-7}$ | 1.44<br>(0.60) | $1.24 \times 10^{-2}$ | 0.78<br>(0.12) | $2.74 \times 10^{-8}$ | +++++ |
| rs75639044 | 7:107,092,631 | *COG5* | C/A | 0.02-0.19 | 0.90<br>(0.18) | $1.20 \times 10^{-7}$ | 1.26<br>(0.78) | $1.04 \times 10^{-1}$ | 0.90<br>(0.18) | $3.63 \times 10^{-8}$ | ----- |
| rs12462756 | 19:50,098,423 | *PRR12* | A/G | 0.17-0.50 | 0.90<br>(0.18) | $4.10 \times 10^{-7}$ | 1.50<br>(0.60) | $1.31 \times 10^{-2}$ | 0.96<br>(0.18) | $3.21 \times 10^{-8}$ | ----- |

Novel genome-wide significant loci discovered in multi-ancestry meta-analysis. Directions of effect were consistent across all five cohorts for all the novel variants (Effect Direction column). The AFR, AHL, EAS, and SAS cohorts were included in the non-EUR meta-analysis (*n* = 37,117).
<sup>a</sup>All lead SNPs are intronic to their 'Nearest Gene,' except for rs7689303, which is approximately 12Mb 3' of *SH3RF1*.
<sup>b</sup>Genotypes of effect and alternate alleles. The minor allele was designated as the effect allele.
<sup>c</sup>EAF (Effect Allele Frequency) range' shows the minimum and maximum effect allele frequencies across the five cohorts.
<sup>d</sup>Effect sizes (Beta) and standard errors (SE) are reported in unit change (minutes of sleep per effect allele copy).

**Figure 3:**
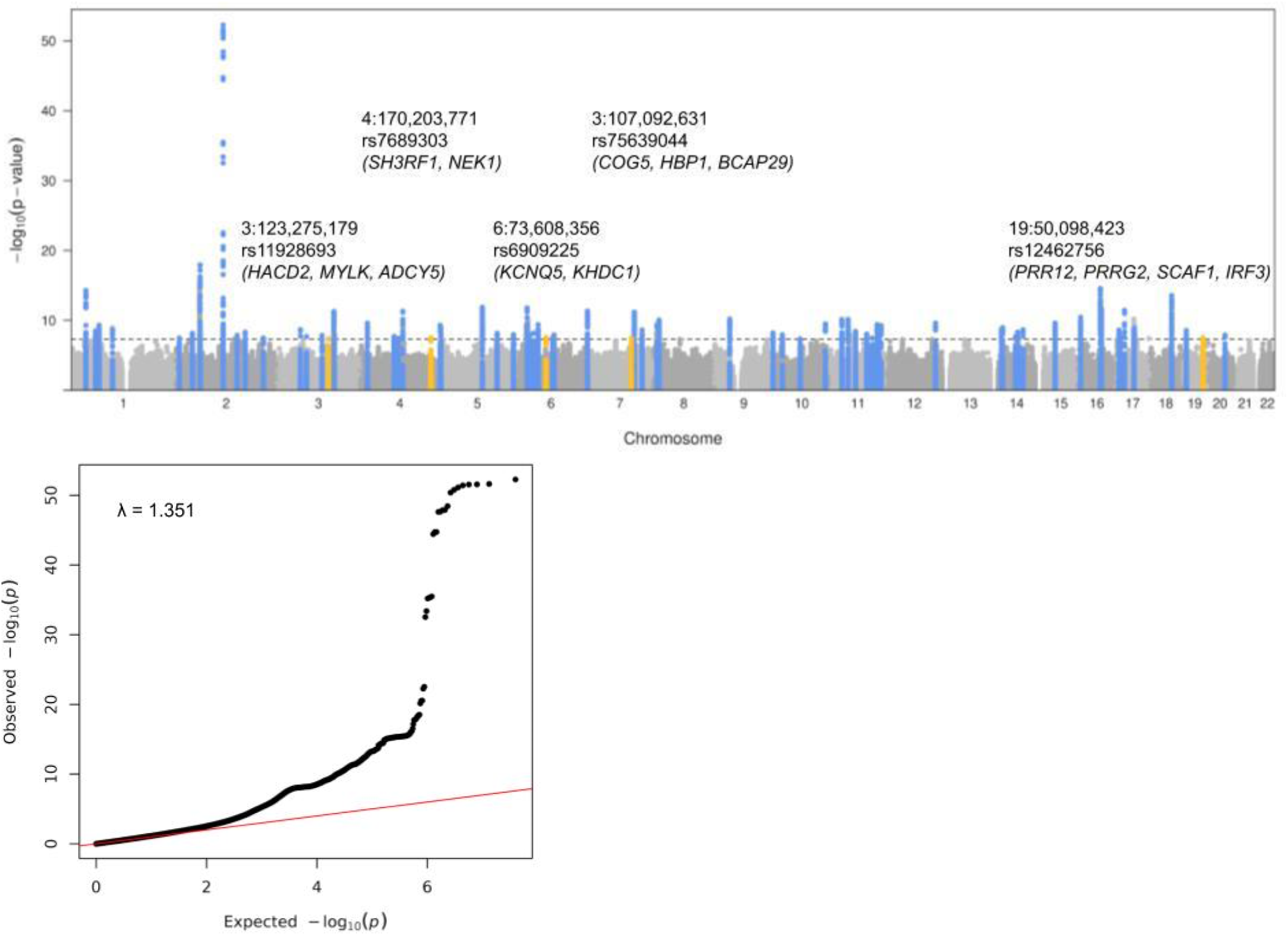
Sleep duration multi-ancestry meta-analysis (n=483,235) Manhattan and Q-Q plots. Sleep duration multi-ancestry meta-analysis (n=483,235) Manhattan and Q-Q plots. The Manhattan plot depicts the five novel loci discovered in the meta-analysis (yellow) and the 68 loci replicated from the European ancestry GWAS (blue). Genomic inflation for the meta-analysis was λ=1.351. This is comparable to inflation values observed for other highly polygenic complex traits, and is also due to the large sample size of the EUR GWAS (λ=1.200) and the meta-analysis structure of the study. Manhattan and Q-Q plots were generated with qqman (R package), and genomic inflation was calculated with GenABEL (Methods).

For the 5 novel loci, replication testing was performed using data from the CHARGE consortium (*n* = 46,810), an independent meta-analysis of 18 GWAS of European ancestry (8). As several of the meta-analysis lead variants were unavailable in CHARGE, we first identified proxy SNPs in strong linkage disequilibrium (all *r^2^* > 0.99 in 1KG Eur population) with the meta-analysis variants (28). For all five proxy variants, directions of effect between CHARGE and the meta-analysis were concordant (binomial p = 3.1 × 10^-2^) (**Table S4**). None of the novel loci were significant in CHARGE after Bonferroni correction, likely due to insufficient power from the smaller cohort size (power to detect associations ≤ 9.0-34.7%).

### Variants at novel loci impact gene expression in the cortex and the cerebellum

To assess the function of the novel loci in influencing gene expression and identify effector transcripts, we examined the eQTLs of those loci in XX brain tissues involved in sleep processes using the GTex cohort (v8) (21) . We selected these tissues as they are highly enriched for expression of genes linked to sleep duration (10). The novel locus represented by lead SNP rs12462756 significantly increases PRR12 gene expression in the cerebellum, cortex, and putamen; it also nominally increases PRR12 expression in the hypothalamus (**Fig. 4a**). Moreover, rs12462756 also significantly increases IRF3 gene expression in the cerebellum (**Fig.4b**). The locus with lead SNP rs75639044 significantly decreases COG5 gene expression in the spinal cord (**Fig. 4c**). The remaining novel loci at or near *SH3RF1*, *KCNQ5*, and *HACD2* did not significantly change the gene expression in any of the brain tissues (**Table S5**).

**Figure 4:**
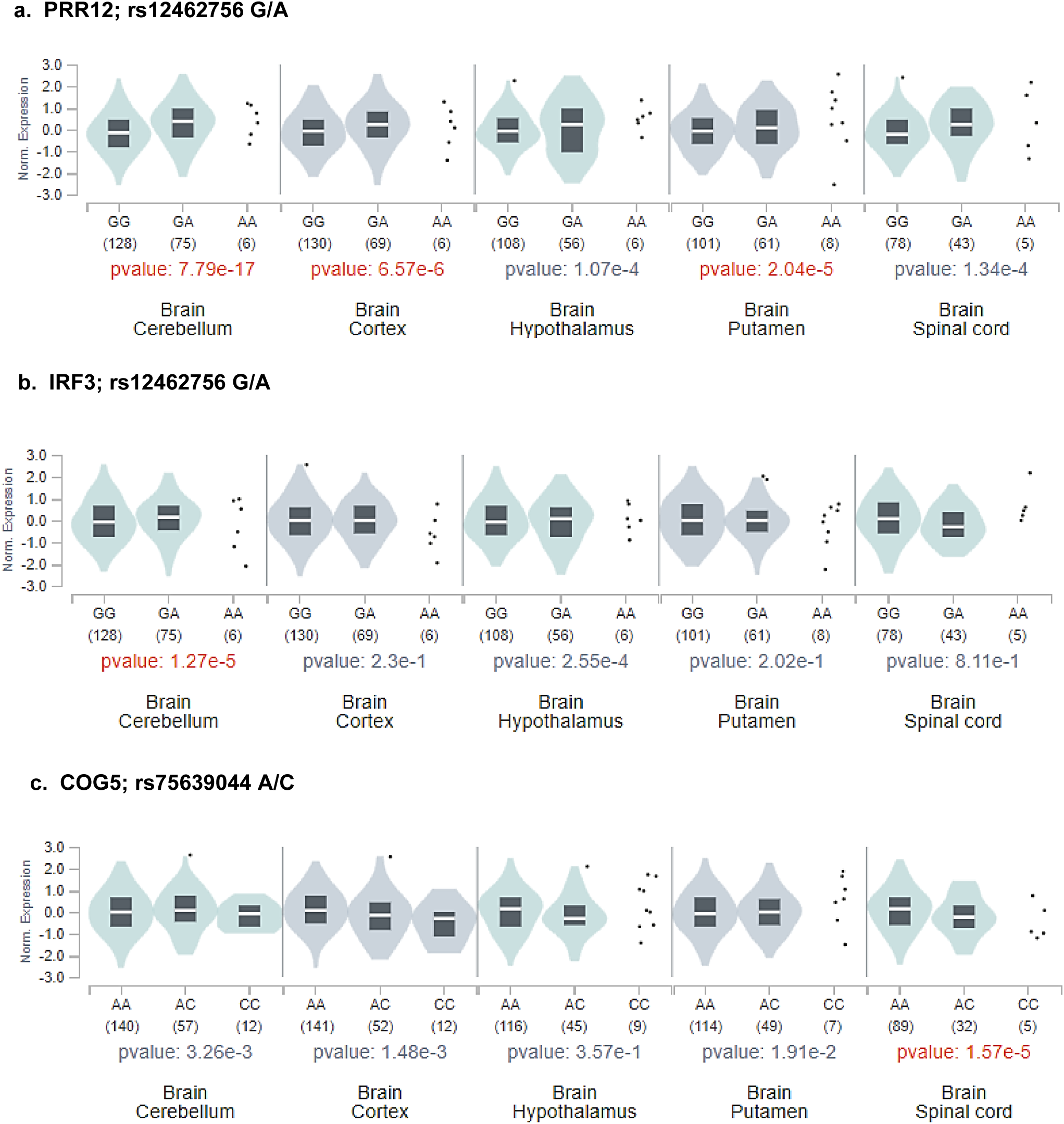
Brain tissue specific eQTLs of novel loci. The eQTLs were examine using GTex V8 resource, and the results are displayed as Normalized gene expression for each locus; the number below box plot is the sample size. **(a)** The sleep duration increasing allele at rs12462756 significantly increase PRR12 gene expression in the cerebellum, cortex, and putamen. **(b)** The locus rs12462756 significantly increase IRF3 gene expression in the cerebellum. The sleep duration increasing allele at rs75639044 significantly decrease COG5 gene expression in the spinal cord.

To distinguish unrelated nearby eqtl signals from eqtl signals driven by the associated loci, we then assessed the colocalization of the genetic association signals and eqtl signals identified above (**Fig. 5; Table S6**) (21, 29). Interestingly, the signals for *COG5*, and *PRR12* colocalized with eQTLs in the cortex, cerebellum, putamen, spinal cord, and hypothalamus. First, the rs12462756 colocalized with *PRR12* expression in the cortex (pp=0.96), cerebellum (pp= 0.99), putamen (pp = 0.83) and hypothalamus (pp=0.73). Second, the rs75639044 signal colocalized (posterior probability (pp) > 0.94) with *COG5* expression in cervical spinal cord. These findings broadly suggest that a common set of brain regions may modulate habitual sleep duration across diverse populations.

**Figure 5:**
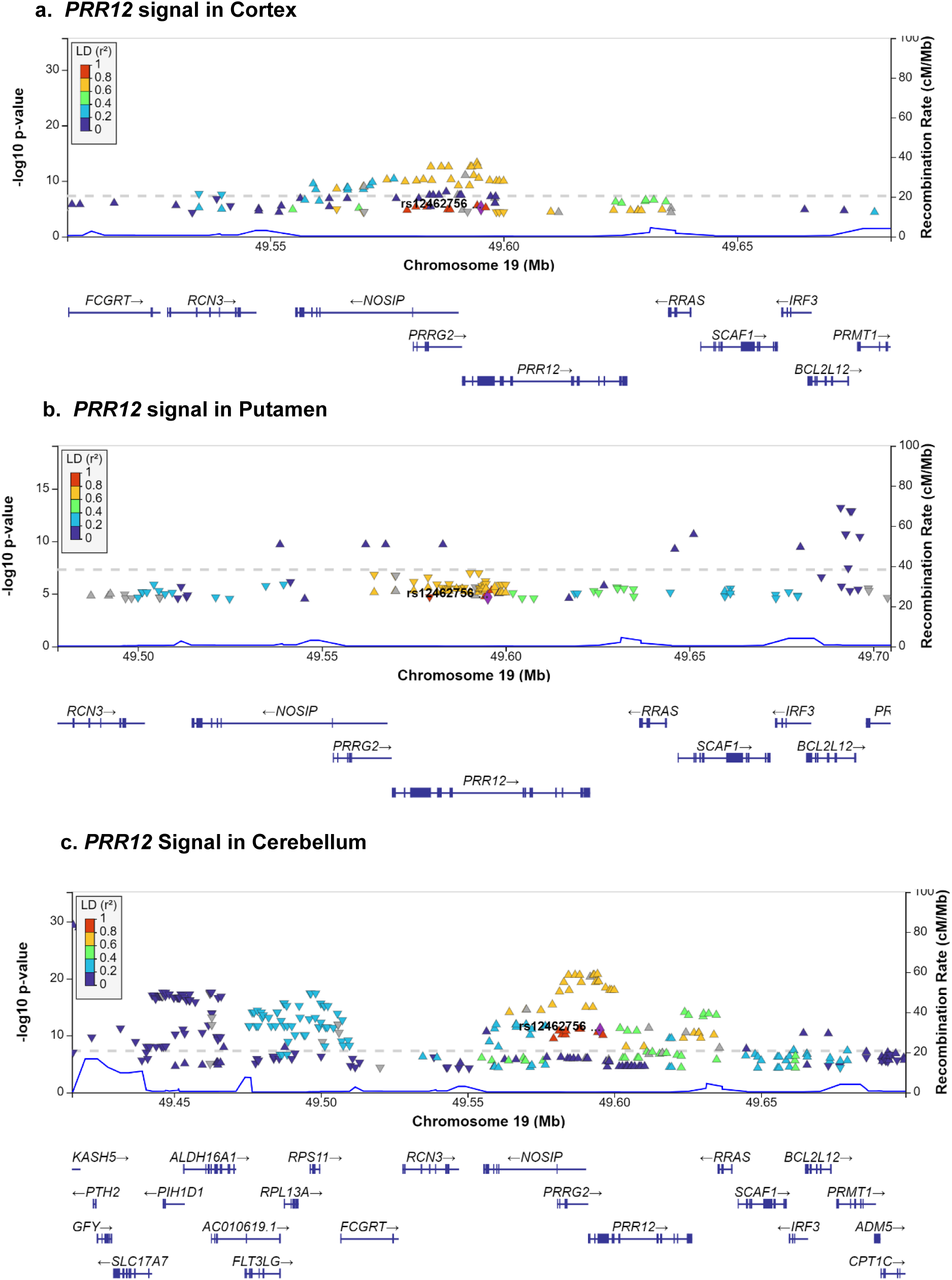
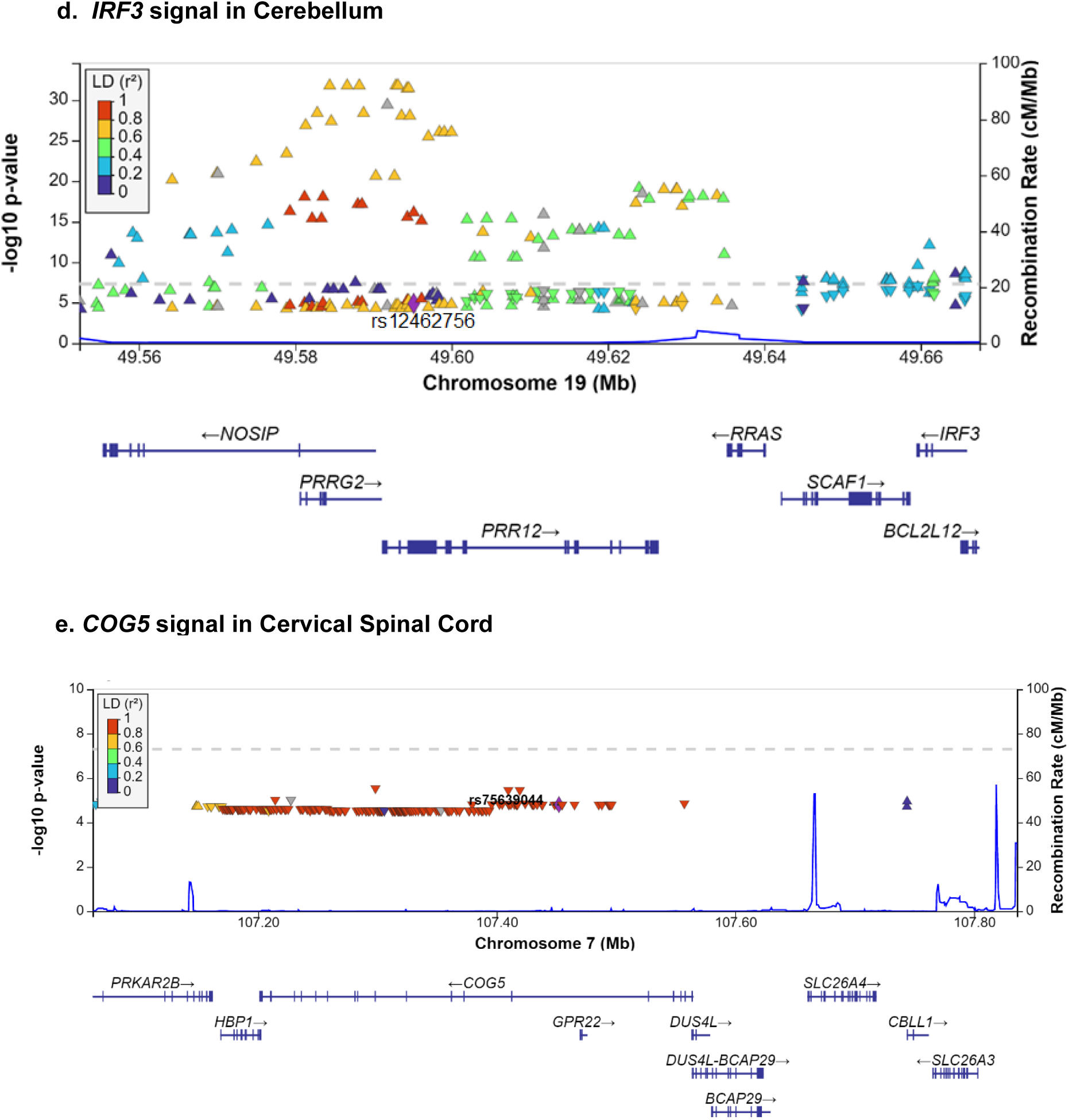
Colocalization of single tissue eQTLs with novel sleep duration loci. Colocalization of single-tissue cis-eQTLs with novel sleep duration loci. Plots of the chromosome 19 locus (*PRR12*) depict the (**a**) The sleep duration increasing allele at rs12462756 significantly increase PRR12 gene expression in the cortex (*n* = 205); (**b**) the putamen (n=170); (**c**) the cerebellum (n=209). (**d**) IRF3 signal in the Cerebellum (n=205). (**e**) The sleep duration increasing allele at rs75639044 significantly decrease COG5 gene expression in the spinal cord (n=126). P-value threshold are in **Table S5**. Lead variants from the cis-eQTL signals were in LD with respective lead variants in the meta-analysis (*r^2^* > 0.8).

### Pleiotropic association to other sleep and disease traits at novel loci

We then examined the genetic associations of the novel signals across six self-reported sleep trait GWAS from the UK Biobank to understand the nature of their impact on regulating sleep (**Table S7**; *P*bonferroni<0.0017) (9,10,30). Three of the novel lead variants are also associated with long (≥9 h per night) sleep duration (*HACD2*, *SH3RF1*, *KCNQ5*, *p* < 9.5 × 10^−5^) while two show nominal association with short (≤6 h per night) sleep duration (*COG5*, *PRR12, p* < 1.3 × 10^−3^). Of these, rs9360616 a variant in LD with the novel rs6909225 in KCNQ5 (r^2^=1 and D’=1) colocalized with long sleep duration (Posterior P = 0.8506; Regional P = 0.9026, Posterior explained by SNP = 0.1242). In a GWAS of napping frequency, three novel signals (*COG5, SH3RF1, PRR12*) showed associations with more frequent napping (*p* < 4.7 × 10^−5^) (31). Of these, rs1978085 (a SNP in strong LD with lead SNP rs75639044 in *COG5* r^2^=1 and D’=1 in 1K EUR population) colocalized with napping frequency (Posterior P = 0.7599; Regional P = 0.9754, Posterior explained by SNP = 0.1197). Nominal associations were observed for morningness (diurnal preference, rs6909225, *p* = 2.8 × 10^−3^), excessive daytime sleepiness (rs75639044, *p* = 1.8 × 10^−3^), and insomnia symptoms (rs12462756, *p* = 7.2 × 10^−3^). These findings suggest that the novel loci may exert pleiotropic effects and that multiple sleep-related traits partially share a common genetic basis.

Next, to identify other traits associated with the novel sleep duration loci, we interrogated a phenome-wide association study (PheWAS) atlas to identify traits associated (causally or pleiotropically) with sleep duration (32). The results highlight novel phenotypes previously unassociated with habitual sleep duration, such as visual acuity and total protein in blood serum (**Table S8**). For example, rs6909225 (*KCNQ5*) is associated with younger age when first started wearing glasses/contacts (**A**/G, *p* = 1.9 × 10^−17^) (32)), and rs12462756 (*PRR12*) is associated with total serum protein (**G**/A, *p* = 6.1 × 10^−10^) (33). The results also highlight traits known to share a genetic basis with sleep duration, including schizophrenia (*r*g=0.26), years of schooling (*r*g=-0.35), and measures of obesity (waist-to-hip ratio, *r*g=0.23) (all *p* < 2.2 × 10^−4^) (10). The effect alleles for the *PRR12* and *SH3RF1* lead variants are associated with longer sleep and increased schizophrenia risk (rs12462756 **G**/A, *p* = 1.2 × 10^−8^ and rs7689303 **G**/T, *p* = 8.7 × 10^−8^) (34), and the effect alleles for the *PRR12* and *COG5* lead variants are associated with increased sleep and reduced cognitive performance (rs12462756 **G**/A, *p* = 1.3 × 10^−6^ and rs75639044 **A**/C, *p* = 8.1 × 10^−6^) (35). Finally, rs11928693 (*HACD2,* 3-hydroxyacyl-CoA dehydratase) is associated with more sleep and increased BMI (p=3.5 × 10^−5^) and body fat percentage (p=4.1 × 10^−6^) (32, 36). These secondary associations may reflect either the same underlying causal variant or gene driving the associations (pleiotropy) or confounding by linkage disequilibrium.

## Discussion

Understanding the genetic basis of sleep duration in diverse ancestry populations can identify biological factors regulating sleep and the molecular links between sleep and disease risks. Furthermore, a robust polygenic score for sleep duration may also be useful to assess links of sleep duration with other diseases in cohorts that may not directly measure sleep (37).

Our first significant finding is that a polygenic score for sleep duration from Europeans is transferable across several diverse ancestry groups. The existence and detection of cross-ancestry genetic transferability depends on multiple factors, including patterns of linkage disequilibrium (whether or not a causal variant is tagged in one or both groups), the frequencies of the lead (tagging) variant, and environmental factors which may affect populations differently. In our first transferability analysis, we found that the sleep duration PGS of 78 EUR loci was significant in 3 of 4 non-EUR GWAS cohorts. The largest effect size was observed in the AFR cohort (0.62 [0.41-0.83] mins/allele), but the effect sizes across all 3 non-EUR cohorts were similar. In our second analysis, we observed that the 10 most significantly associated EUR variants were highly transferable to the non-European GWAS cohorts, and, as expected, rare minor alleles were associated with lower transferability. At *PAX8*, *FTO*, and *PPP2R3A*, the minor alleles of lead variants were relatively common, and their effect sizes and directions were highly conserved across cohorts. At *VRK2* and *BANK1*, minor alleles of lead variants were rarer and showed limited transferability. Finally, we observed that multiple genome-wide PGS were significant in AFR and SAS at multiple p-value thresholds. This suggests that genetic variation associated with habitual sleep duration is distributed across the genome. This is consistent with prior findings, as meta-analyses of multiple complex traits have found that >0.1% of all SNPs may be causal for highly polygenic complex traits (32). Overall, our results suggest that the rich genetic architecture of habitual sleep duration is at least partially transferable across divergent ancestry groups.

Interestingly, the PGS composed of 78 EUR variants was not significant in the AHL cohort, which could arise from differences in participant demographic, behavioral, phenotypic and/or genetic characteristics between UKBB and the HCHS/SOL cohort (38). First, the AHL GWAS cohort was younger (mean age 46.2 (14.1)) and more female (60.2%) than the UKBB cohort (mean age 57.3 (8.0), 54.7% female). Given these factors, AHL participants may have had greater responsibilities related to childcare, which could have impacted their sleep schedules and biased self-report of wake and bed times. Second, many HCHS/SOL participants reported working shifts. While shift workers were excluded from the AHL GWAS study to limit confounding from atypical sleep schedules, participants who work multiple jobs or irregular hours were not excluded. This may have led to a bias toward overestimating sleep time. Third, sleep duration was calculated in HCHS/SOL based on self-reported bed and wake times and validated by actigraphy in a subset of the population (r=0.48) (39), whereas UKBB and J-MICC participants directly reported their sleep durations in more imprecise 1 h increments. These two measures yield correlated (r = 0.4-0.7) (24, 39)but distinct estimates of sleep duration, which could also have affected our estimate of transferability. Fourth, genetic differences between European and Hispanic/Latino ancestry groups could underlie observations of transferability. Future studies are needed to better understand the genetic architecture of sleep duration in Admixed Hispanic/Latino groups.

Importantly, multi-ancestry meta-analysis of self-reported sleep duration identified 5 novel and 68 known genetic loci associated with self-reported sleep duration. We found that these novel loci are associated with other sleep-related traits, schizophrenia risk, years of schooling, and BMI. Many of these traits are known to share a genetic basis with sleep duration in European populations (10) and our findings indicate that non-European populations may have similar patterns of genetic architecture. Moreover, in our colocalization analysis, the 7 out of the 13 tested brain tissues were overrepresented. This suggests that brain regions known to influence sleep (frontal cortex, anterior cingulate, and nucleus accumbens) as well as some that have received less attention such as the cerebellum (40), are key regulators of habitual sleep duration in cohorts of diverse ancestry.

Results from our colocalization analysis shed light on a potentially causal mechanism at the novel *PRR12* locus (Proline Rich 12) (chr12: rs12462756). We found that the rs12462756 signal colocalized with the *PRR12* eQTL in the cortex, cerebellum, putamen, and hypothalamus (**Fig. 5** and **Table S6**), and that higher *PRR12* expression was associated with napping. Prior work reported *PRR12* to be involved in inflammatory disorders (41, 42); and to the best of our knowledge, we are the first to report on the implication of *PRR12* in both sleep duration and napping. These findings suggest that genetic variation in *PRR12* may increase habitual sleep duration via the cortex and the cerebellum. As with other associations, *in vivo* analyses will be needed to fully establish the mechanism of action through which *PRR12* influences sleep duration.

Links between the novel loci and other traits underscore the deleterious connections between long and short habitual sleep duration and cardio-metabolic and psychiatric disorders. First, rs11928693 (*HACD2*, 3-hydroxyacyl-CoA dehydratase) is associated with longer sleep duration, increased BMI, and increased body fat. While short sleep is widely understood to be linked to obesity (42–44), this finding suggests long sleep may also be connected to obesity. Second, the lead variants at the novel *PRR12* (rs12462756) and *COG5* (rs75639044) loci are associated with longer sleep and reduced cognitive performance (37). These associations add to evidence that genetic factors for both long and short sleep duration are associated with lower educational attainment (10). Finally, the lead variants of the *PRR12* (rs12462756) and *SH3RF1* (rs7689303) loci are associated with longer sleep and increased schizophrenia risk. Unlike obesity and educational attainment, schizophrenia risk has been found to share a genetic basis with long sleep duration but not short sleep duration (43, 44). Further colocalization analyses are needed to show that these associations are not due to separate signals, and ultimately, in vitro studies will be needed to unravel the biological mechanisms that underpin these trait effects.

The strengths of our study include the ethnic diversity of its cohorts, the assessment of transferability at multiple genomic levels (individual alleles, PGS of lead variants, and genome-wide PGS) and the use of colocalization analyses to examine variant impact on gene expression and pleiotropy. The cross-ancestry transferability of a polygenic score for sleep duration demonstrates shared genetic architecture of sleep duration across ethnic groups. The diversity of our meta-analysis aided the detection of novel loci, as cohort differences in allele frequency, physiologic effect, and evolutionary history add discovery power (20). While the non-EUR, single-ancestry cohorts were underpowered for secondary analyses, we expanded upon our primary findings by identifying pleiotropic associations for the novel loci and conducting a colocalization analysis.

However, our study has some limitations. First, this study was not designed to rigorously evaluate the genetic transferability at the single variant and gene level; the sample sizes of the non-European cohorts were too small to assess the transferability of individual genetic variants and loci. In addition, this study did not assess whether associations between the 5 novel sleep duration loci and other traits (in our PheWAS analysis) may be due to distinct genetic signals or pleiotropy. Last, the assessment of transferability to Admixed Hispanic/Latino groups may have been limited by demographic and phenotyping differences between the EUR cohort and AHL cohorts. Given these limitations, we advocate for increased recruitment of non-European participants to biobank studies. Greater representation of non-European populations in biobanks will improve the detection of rare variants, aid the creation of ancestry-specific PGS, and facilitate the equitable translation of scientific advances from genomics to multiple worldwide populations.

In summary, we found that the genetic basis of habitual sleep duration in European ancestry groups partially transfers to non-European ancestries. Moreover, a common set of phenotypes (schizophrenia, BMI, etc) and brain regions (cortex, cerebellum and hypothalamus) are likely involved in maintaining habitual sleep duration in multiple groups of divergent ancestry. While larger population-specific GWAS will be necessary to detect rare variants associated with sleep duration, we show that 78 variants from EUR strongly associate with sleep duration in multiple non-European ancestry cohorts. Overall, this study advances the understanding of the genetic architecture of habitual sleep duration and contributes to the improved detection and treatment of sleep diseases and related neuropsychiatric disorders.

## Methods

### Cohort selection and design

We included study participants from three cohort studies, including the UK Biobank Study (45), the Hispanic Community Health Study/Study of Latinos (HCHS/SOL) (46), and the Japan Multi-Institutional Collaborative Cohort Study (J-MICC) (47). The UK Biobank (UKBB) is a study of 500,000 UK volunteers which was designed to investigate the environmental and genetic determinants of human traits and diseases. The European ancestry cohort (EUR, *n* = 446,118) was subset from the UKBB, and this cohort comprised the largest published GWAS of self-reported sleep duration to date. We also identified 7,518 African ancestry (AFR) and 7,660 South Asian ancestry (SAS) participants from the UKBB for their respective GWAS cohorts. HCHS/SOL is a community-based prospective cohort study of 16,415 self-identified Hispanic/Latino persons aged 18-74 years at screening to assess the role of acculturation in the prevalence and development of disease, and to identify factors playing a protective or harmful role in the health of Hispanics/Latinos. Based on phenotyping and genotyping inclusion criteria (**Table S2**) we selected 8,726 participants for the Admixed Hispanic/Latino (AHL) GWAS (48). Lastly, GWAS for 13,618 Japanese participants from J-MICC was conducted by Nishiyama et al and analyzed here as the East Asian ancestry (EAS) cohort (11). Participants provided informed consent in each study according to institution specific protocols.

### Sleep duration phenotyping and covariate measures

Sleep duration phenotyping is summarized in **Table S2**. Briefly, in the UK Biobank, study participants (n ∼ 500,000) self-reported sleep duration at baseline assessment. Participants were asked: About how many hours sleep do you get in every 24 h?, with responses in hour increments. Extreme responses of less than 3 h or more than 18 h were excluded and ‘Do not know’ or ‘Prefer not to answer’ responses were set to missing. Participants who self-reported any sleep medication were excluded. In HCHS/SOL AHL, 12,803 participants with high quality genetic data reported habitual weekday and weekend bed and wake times. Shift workers, participants with sleep duration < 3 h or > 14 h on weekdays or weekends, and those on sleep medications were excluded (n=4,077). Average sleep duration was calculated as a weighted average (5/7 weekday + 2/7 weekend) of difference between self-reported bedtime and wake time for the remaining participants. In the J-MICC, as previously described (11), subjects reported usual sleep duration in numbers (integer and half-integer values), and respondents with self-reported sleep medication use were excluded.

### Genotyping and quality control

The genotyping and quality control characteristics of this study’s cohorts are briefly described here; for additional information see the corresponding biobank publications (46,47,49) and the genotyping and phenotyping summaries by cohort (**Table S2)**. Genotyping in the UK Biobank was conducted with the UK BiLeVe and UK Biobank Axiom arrays. Samples with sex chromosome abnormalities or high missingness or heterozygosity rates were removed, and 488,377 samples were available for study at the time of this analysis. Marker-based QC tested for batch effects, plate effects, Hardy– Weinberg equilibrium, sex effects, array effects, and discordance across control replicates. In this study, we selected the AFR and SAS GWAS cohorts using principal components of ancestry and self-reported ethnicity (Methods, **Table S1**).

In HCHS/SOL, subjects were genotyped with a custom array which consisted of a HumanOmni2.5-8 v1.1 BeadChip array with ∼150,000 ancestry-informative markers and hits from previous GWAS (46, 48). Samples with inconsistent sex, gross chromosomal anomalies, or contamination were excluded (50), as were variants with high missingness, Mendelian errors, duplicate-sample discordance, or deviations from Hardy-Weinberg equilibrium. In J-MICC, subjects were genotyped with the HumanOmniExpressExome-8 v1.2 BeadChip (47) and samples with inconsistent sex, relatedness (using PLINK pairwise identity-by-descent estimation (51), or non-Japanese genetic ancestry (as assessed with principal components of ancestry (52) were removed. SNPs with deviations from known MAF values or Hardy-Weinberg equilibrium were removed, as were variants with low call rates.

### Selection of AFR and SAS cohorts from UKBB

We selected the AFR and SAS GWAS cohorts on the basis of principal components of ancestry (PCAs) and self-reported ethnicity. First, we generated 40 PCAs for all UKBB participants and subset the 486,065 participants into four ancestry clusters using K-means clustering (**Table S1**). The four clusters approximately identified the four major ancestry designations of the UKBB (“European”, “Asian”, “African”, and “Mixed, other”). From the African ancestry cluster (n = 9,039), we then subset 7,518 participants who self-identified as “Black or Black British”, “Caribbean”, “African”, or “Any other Black background” for the African ancestry GWAS cohort (AFR). Similarly, from the South Asian ancestry cluster (n = 13,249), we subset 7,660 participants who self-identified as “Asian or Asian British”, “Indian”, “Pakistani”, or “Bangladeshi” for the South Asian ancestry GWAS cohort (SAS).

To validate the AFR and SAS cohorts, we assessed their similarity to Human Genome Diversity Panel (HGDP) (57) reference populations with TRACE (fasT and Robust Ancestry Coordinate Estimation) (58). Plots depicting the first four PCAs for the six major HGDP reference regions (“Africa”, “Europe”, “Middle East”, “C/S Asia”, “East Asia”, “Oceania”, “America”) were created, and subjects from the AFR and SAS cohorts superimposed (**Figure S1**). We assessed the AFR cohort for overlap with the ‘African’ HGDP reference population and the SAS cohort for overlap with the ‘C/S Asia’ HGDP reference population. The EUR cohort (n = 446,118) was previously selected by Dashti et al using 40 PCAs (10).

### Genome-wide association analyses

We conducted genome-wide association analyses (GWAS) for the AFR and SAS cohorts and obtained GWAS summary statistics for the EAS, EUR, and AHL cohorts. For the AFR (n=7,288) and SAS (n=7,485) cohorts, we used PLINK v1.9 (53) to generate linkage disequilibrium (LD) score files and conducted GWAS with BOLT-LMM v2.3. (54). Covariate corrections for genotyping array, sex, age, 10 PCs of ancestry, and a genetic correlation matrix were made for the AFR and SAS GWAS. Furthermore, for these two cohorts heritability (*h^2^*) was calculated with GCTA-GREML(22), and we excluded variants with MAF < 0.01 in this analysis.

For the HCHS/SOL cohort (AHL, n=8,726), GWAS analysis was performed as described by Conomos et al. (48) ; specifically, a mixed model analysis was conducted using the GENESIS R package accounting for population stratification by adjusting for 5 PCs of ancestry, and for recent relatedness using a kinship matrix (55). To account for HCHS/SOL study design and environmental sources of correlations, correlation matrices were used to account for household and block unit sharing among participants, and adjusted for log of sampling weights. Models were further adjusted for sex, age, study center, and Hispanic genetic background group (**Table S2**). Heritability was calculated as the proportion between the estimated variance component corresponding to the kinship matrix, and the sum of all estimated variance components (household sharing, unit sharing, kinship, and residual variance). Variance components were estimated from a linear mixed model implemented in the GENESIS R package, with the same covariates as the main analysis. Nishiyama et al conducted the GWAS and secondary analyses for the J-MICC cohort (EAS, n=13,618) (11). SNPTEST with the ‘expected’ method was used for the association analysis (56), and sleep duration was regressed on age, sex, the first 5 PCs of ancestry, and allele dosage from imputation (**Table S2**). Heritability was calculated with GCTA-GREML (22).

For all GWAS cohorts (AFR, AHL, EAS, EUR, and SAS), we calculated genomic inflation factors (λ) with the R package GenABEL (57). The inflation factors (1.01 - 1.43) were comparable to previously reported values for other highly polygenic complex traits. Lastly, prior to conducting the multi-ancestry meta-analysis, we excluded variants with minor allele frequencies (MAF) < 0.01 or imputation scores (INFO) < 0.8 from each GWAS cohort.

### Polygenic Score (PGS) using EUR sleep duration signals

In our examination of cross-ancestry transferability, we first assessed the association of the PGS comprised of 78 EUR variants with habitual sleep duration in each of the non-European GWAS cohorts. First, we identified lead variants from the 78 significant loci (*P* < 5 × 10^−8^) in the EUR GWAS and subset these variants from the single-ancestry GWAS.

We then normalized the 78 SNPs’ effect sizes in the EUR GWAS (to a mean of 1.0 min/allele). The R package GTX (Genetics ToolboX) was used this analysis (23).Of the 78 SNPs from EUR, the EAS GWAS included 61, the AHL GWAS included 75, and the AFR and SAS GWAS each included 77 SNPs. We did not include any proxy or alternate SNPs in the PGS calculations. To create the cross-ancestry transferability plot (**Fig. 1**), we identified the 10 most significant loci in the EUR GWAS and then identified the lead variants for these loci in our meta-analysis (**Table S3**). Minor allele frequency data was from the GWAS cohorts, and the plot was then generated in Excel 2016. Finally, we implemented PRSice v2 to assess the aggregate burden of sleep duration associations from the EUR cohort to the AFR and SAS ancestry cohorts (25). PRSice analyzed the summary statistics from the base cohort (EUR) and individual-level genotyping and phenotyping data from the target cohorts (AFR and SAS).

### Multi-ancestry meta-analyses

We then ran a multi-ancestry meta-analysis (all five cohorts) and a non-EUR meta-analysis (four non-EUR cohorts) using METAL (26). The multi-ancestry meta-analysis (*n* = 483,235) analyzed 19,794,469 SNPs from five GWAS in a fixed-effects model. 7,944,585 SNPs were present in four or more GWAS and 5,233,370 variants were present in all five GWAS (**Table S3**). We identified five novel loci in the multi-ancestry meta-analysis and generated regional association plots using LocusZoom (**Fig. S3**) (58). We also conducted a fixed-effects meta-analysis which included the four non-European ancestry cohorts (*n* = 37,117). Here, 8,188,301 variants were present in ≥ 3 cohorts and 6,014,571 variants were present in all 4 cohorts. For both meta-analyses, we generated Manhattan and QQ plots with the R package qqman (59).

### Replication testing in CHARGE

We utilized CHARGE consortium data for replication testing of the 5 novel sleep duration loci (**Table S4**) (8). As the five lead SNPs were unavailable in CHARGE, we chose proxy SNPs (all *R^2^* > 0.99) using LDProxy (28). We also calculated the detection power at p=0.01 (Bonferroni correction threshold) for each of the novel variants in CHARGE using Quanto v1.2 (60). We assumed a MAF=0.10 and an additive genetic model.

### Identifying pleiotropic and sleep trait associations for novel loci

To gain a greater understanding of the impact of the 5 novel loci on sleep-related and pleiotropic traits and disorders (**Table S7**), we first examined associations between the lead variants for the novel loci in GWAS of long sleep duration (≥9 h per night), short sleep duration (≤6 h per night), napping frequency (10), daytime sleepiness, insomnia (31) and morningness (diurnal preference) (9). For diurnal preference, participants were prompted to answer the question “Do you consider yourself to be?” with one of six possible answers: “Definitely a ‘morning’ person”, “More a ‘morning’ than ‘evening’ person”, “More an ‘evening’ than a ‘morning’ person”, “Definitely an ‘evening’ person”, “Do not know” or “Prefer not to answer”. To identify secondary associations for the novel loci, we then conducted a phenome-wide analysis (PheWAS) using the CTGLab’s GWAS Atlas web interface and report associations with *p* < 5.0 × 10^−5^ (**Table S8**) (32). **Colocalizing novel loci with single-tissue cis-eQTLs**

For the novel loci, we first designated the primary gene at each locus as the putatively causal gene which would be tested for signal colocalization with eQTLs from the brain (**Table S3**). We selected 13 brain tissues and pituitary tissue from GTEx as these tissues were found to be highly enriched regions for gene expression signals related to habitual sleep duration (**Table S5**) (10). We then assessed for colocalization between the brain eQTLs and and the loci from the meta-analysis with HyPrColoc (Hypothesis Prioritisation in Multi-Trait Colocalization), and used HyPrColoc’s default parameters (prior.2=0.98, bb.alg=F) (**Table S6**) (29). HyPrColoc reports the posterior probability of colocalization (pp), the regional posterior probability of colocalization, a candidate causal SNP, and the percent contribution of the candidate SNP to the total posterior probability. We generated regional association plots for three loci (*PRR12*, *IRF3* and *COG5*) with LocusZoom (58).

## Author contributions

This study was designed by H.S.D, J.M.L. and R.S. All co-authors participated in the acquisition, analysis, and/or interpretation of data. T.S., R.S., S.R., and P.Z. conducted GWAS for the HCHS/SOL cohort, T.N., M.N., Y.M., K.W., and S.S. conducted GWAS for the J-MICC cohort, and B.H.S., Y.S., J.M.L., H.S.D and R.S. performed remaining analyses. B.H.S and R.S. wrote the manuscript and all co-authors reviewed and edited the manuscript before approving its submission. R.S. is the guarantor of the work and, as such, had full access to all the data in the study and takes responsibility for the integrity of the data and the accuracy of the data analysis.

## Acknowledgements

This research has been conducted using the UK Biobank Resource (UK Biobank application number 6818). We would like to thank the participants and researchers from the UK Biobank study, J-MICC, and HCHS/SOL who contributed or collected data. H.S.D. and R.S. are supported by NIH R01DK107859, NIH R01 DK102696 (R.S.) and MGH Research Scholar Fund (R.S.). J.M.L. is supported by NIH/NHLBI K01 HL136884. H.M.O. was supported by Instrumentarium Science Foundation, Yrjö Jahnsson Foundation and Academy of Finland (#269517). T.S was partially supported by and SR was supported by NHLBI R35 HL135818. T.N., M.N., Y.M., K.W., and S.S were supported by Grants-in-Aid for Scientific Research for Priority Areas of Cancer (No. 17015018) and Innovative Areas (No. 221S0001) Japan Society for the Promotion of Science (JSPS) KAKENHI Grant (No. 16H06277 [CoBiA]) from the Japanese Ministry of Education, Culture, Sports, Science and Technology. T.N., M.N., Y.M., K.W., and S.S were also supported in part by funding for the BioBank Japan Project from the Japan Agency for Medical Research and Development since April 2015, and the Ministry of Education, Culture, Sports, Science and Technology from April 2003 to March 2015.

## Supp. Figures

**Figure S1.**
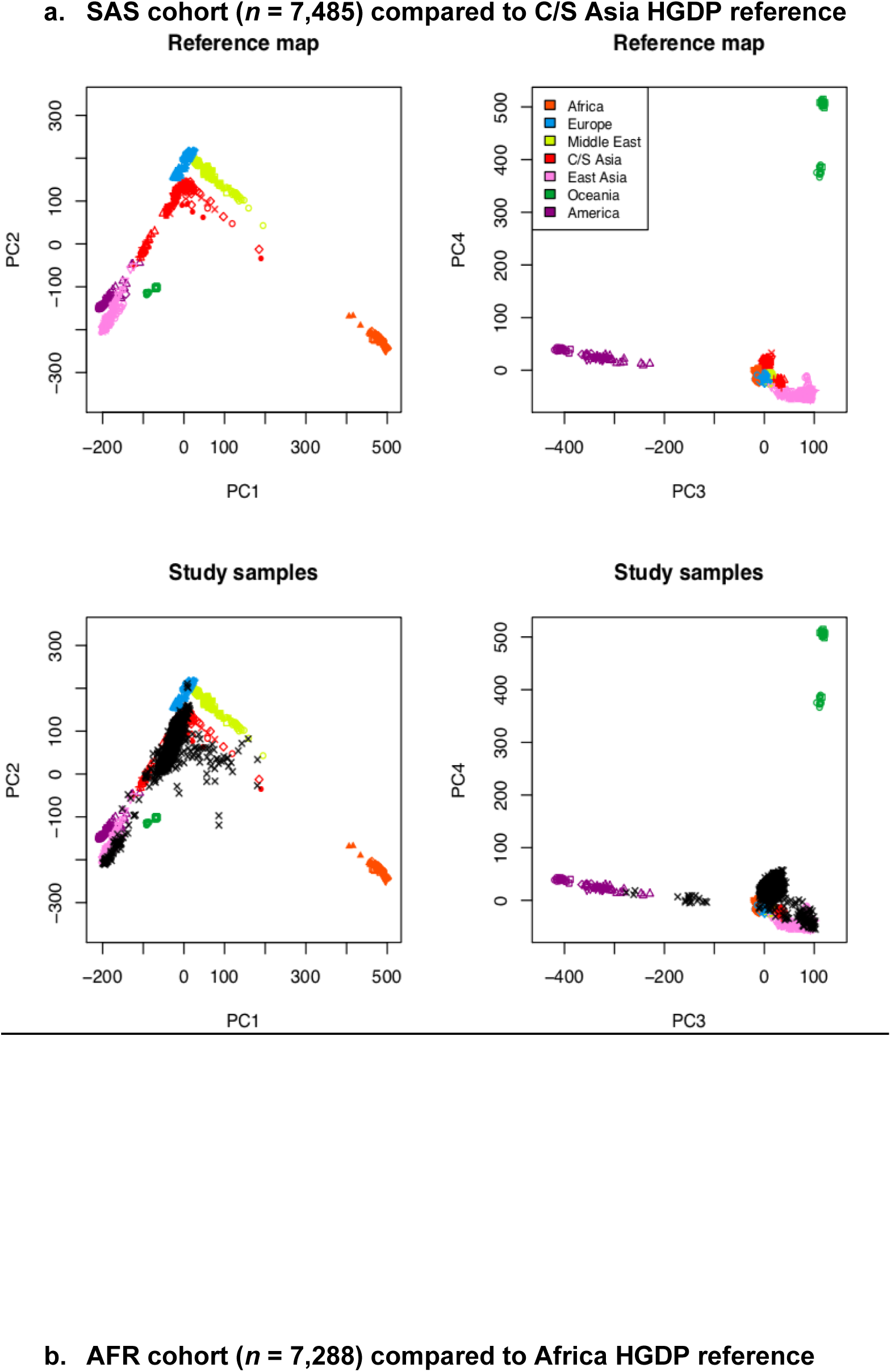
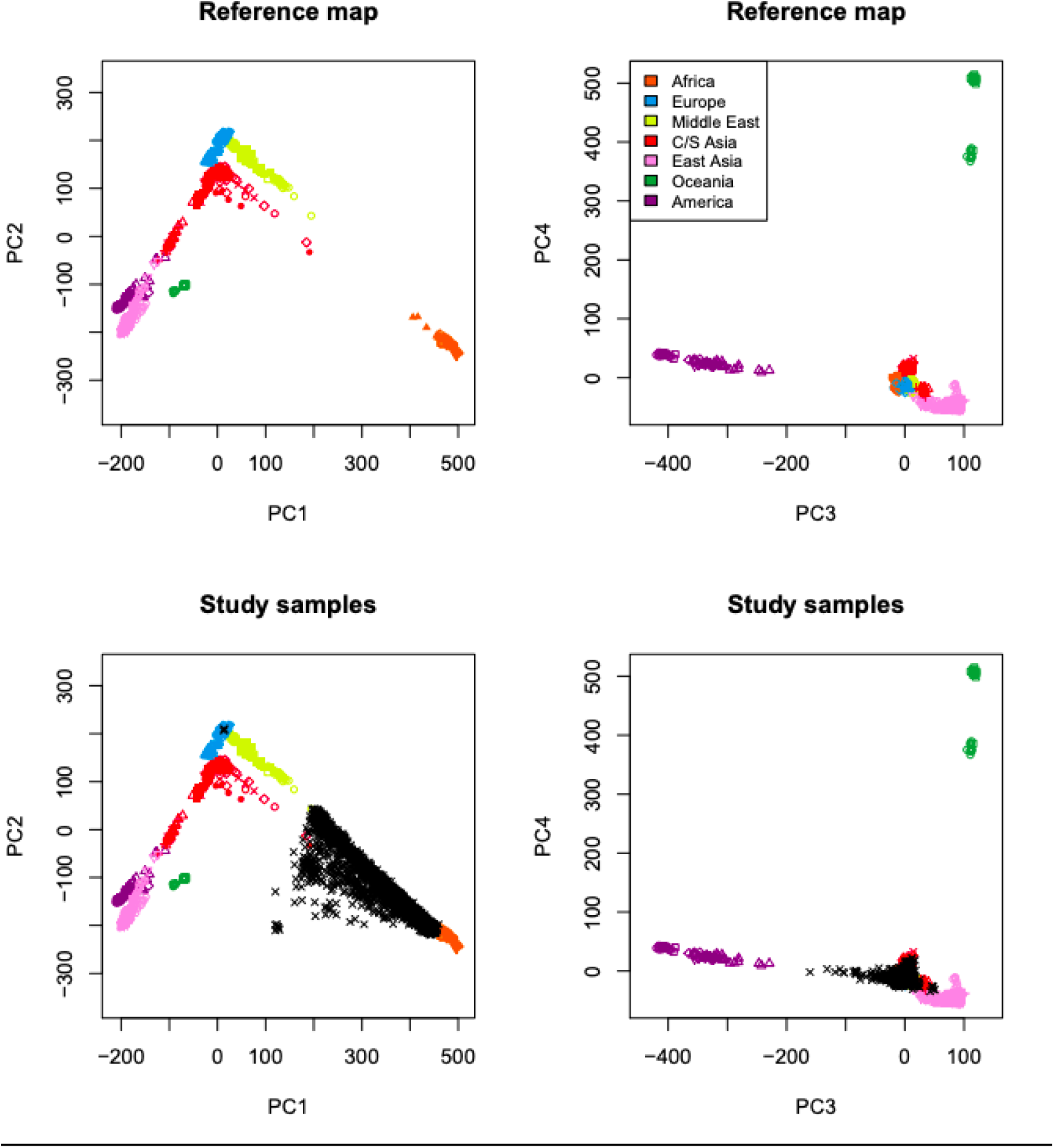
SAS and AFR cohorts (UKBB) validated with HGDP ancestry references. a. SAS cohort (*n* = 7,485) compared to C/S Asia HGDP reference b. AFR cohort (*n* = 7,288) compared to Africa HGDP reference SAS and AFR cohorts (UKBB) validated with HGDP ancestry references. To construct the South Asian ancestry (SAS) GWAS cohort, a group of 13,630 genetically similar individuals were identified in a k-means clustering (k=4) analysis. From this cluster, 7,660 individuals who self-identified as Indian, Pakistani, Bangladeshi, Asian or Asian British were subset for SAS. For the African ancestry (AFR) GWAS cohort, a group 7,518 individuals identifying as Black or Black British, Caribbean, African, or Any other Black background were selected from a second cluster of 9,280 individuals. TRACE (fasT and Robust Ancestry Coordinate Estimation) was used to generate ancestry plots.

**Figure S2.**
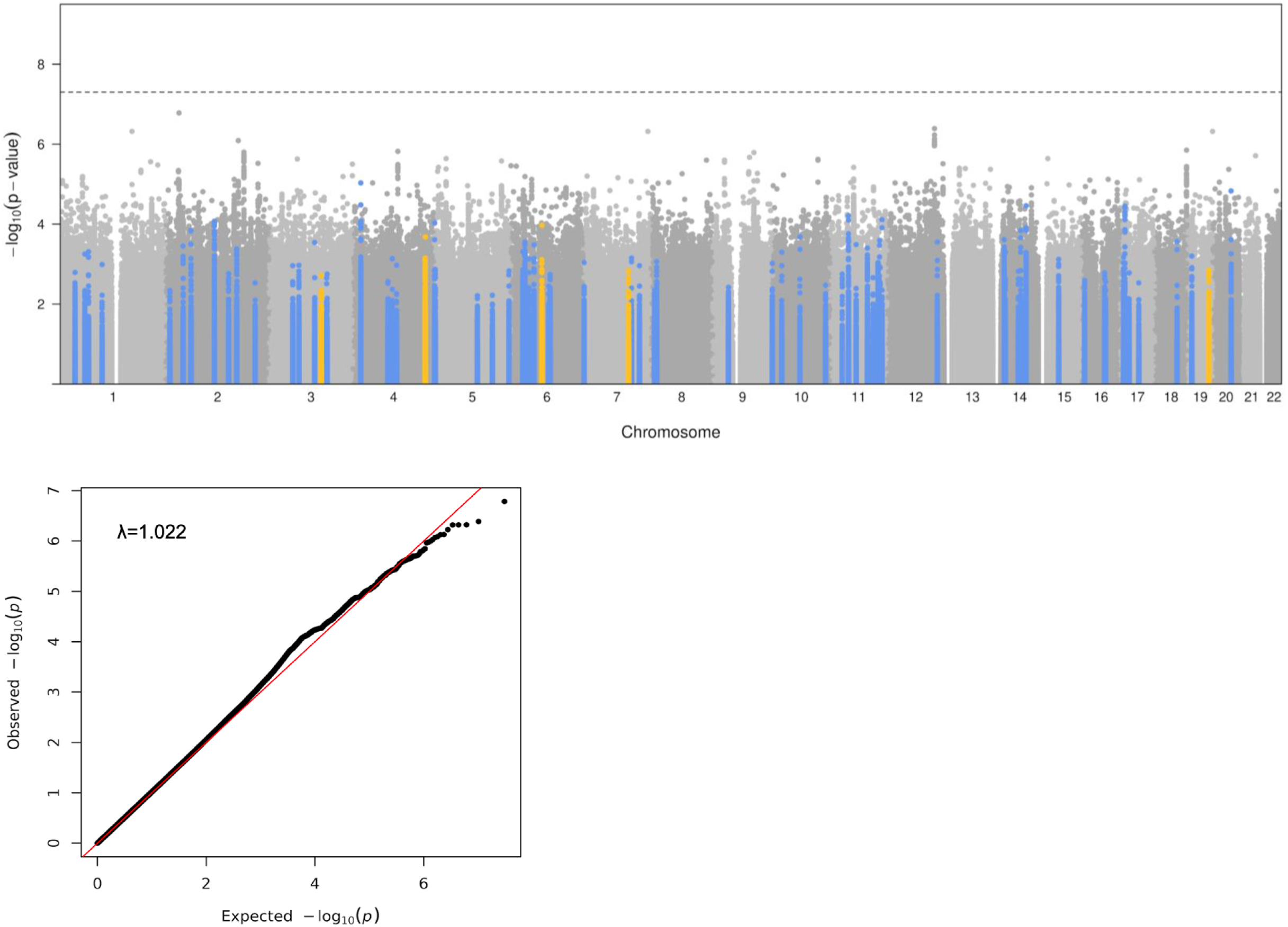
Manhattan and Q-Q plots for non-EUR meta-analysis (*n* = 37,117) Manhattan and Q-Q plots for non-EUR meta-analysis (n = 37,117). The Manhattan plot depicts the 68 loci (blue) that were replicated in the multi-ancestry meta-analysis (n = 483,235) and the five novel loci (yellow) discovered in the meta-analysis. No genetic loci were significant (*p* < 5.0 × 10^−8^) in the non-EUR meta-analysis. Plots were generated with qqman (R package), and genomic inflation (λ=1.022) was calculated with GenABEL (Methods).

**Figure S3.**
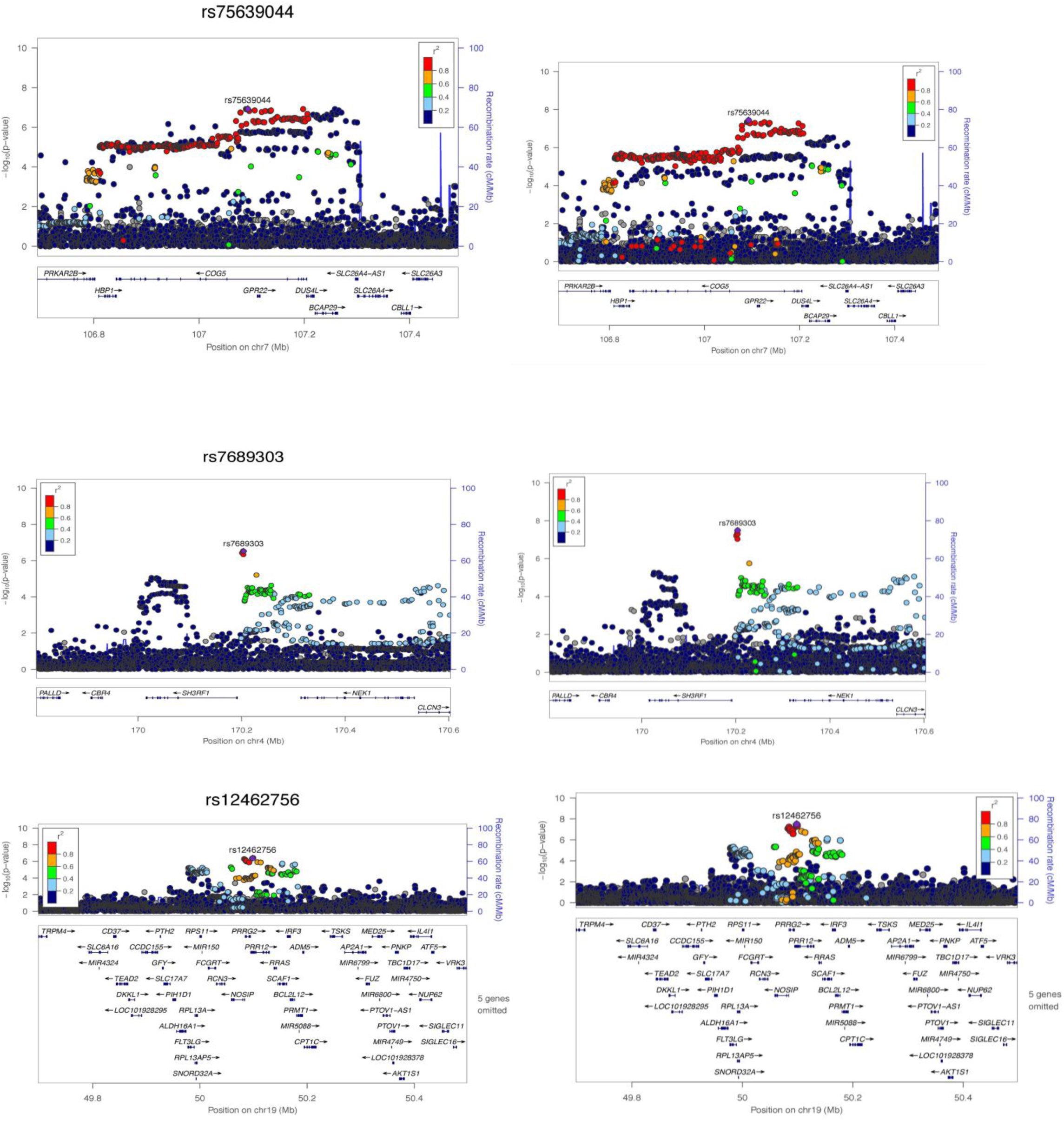
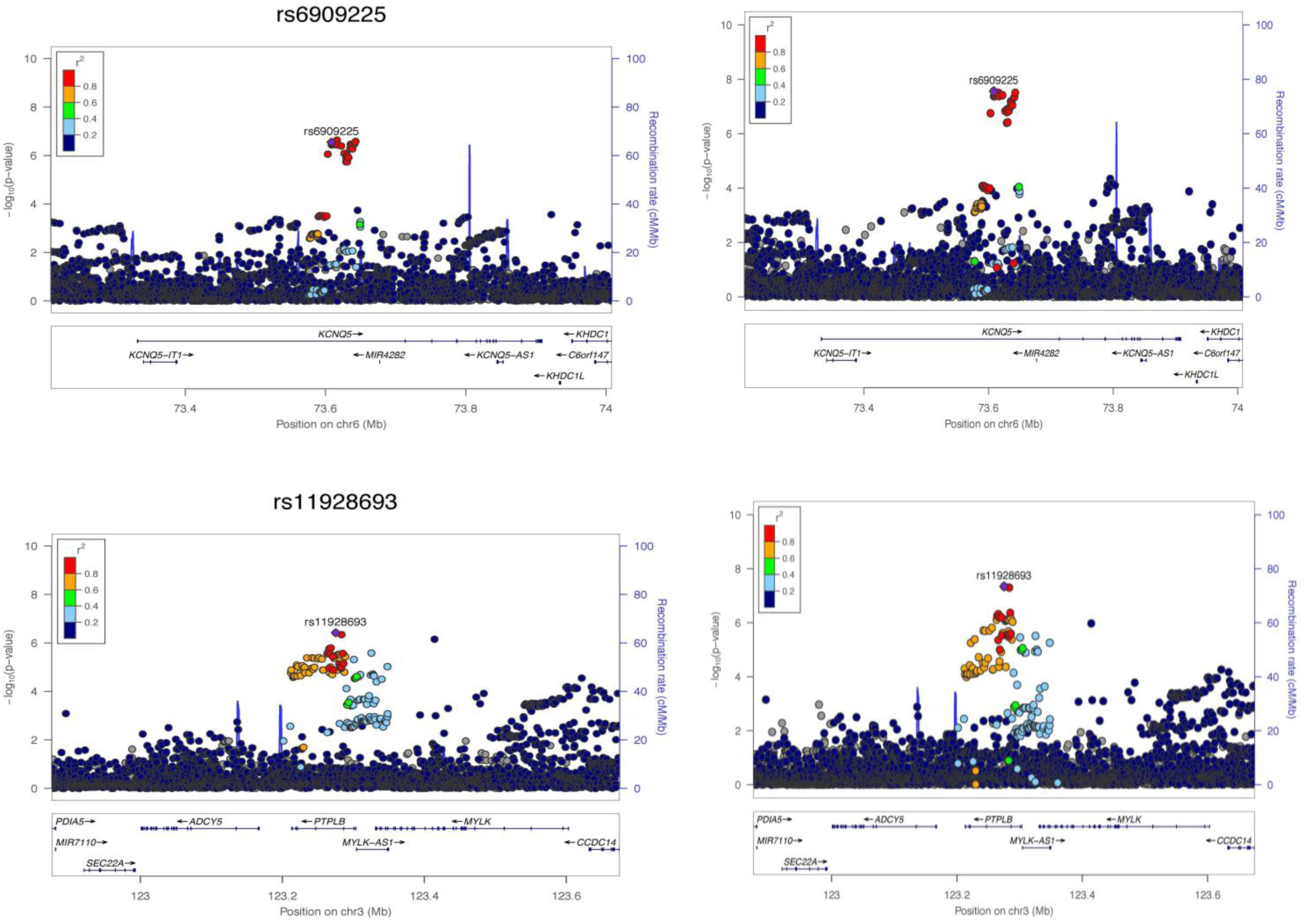
Five novel loci in EUR ancestry GWAS (n = 446,118) and multi-ancestry meta-analysis (n = 483,235) Five novel loci in EUR ancestry GWAS (n=446,118) and multi-ancestry meta-analysis (n=483,235). The novel loci (on chromosomes 3, 4, 6, 7, and 19) are shown. Plots are centered on lead variants with 400 kb upstream and downstream from the lead variant. European ancestry LD was chosen as the LD reference for all plots (Methods).

**Figure S4.**
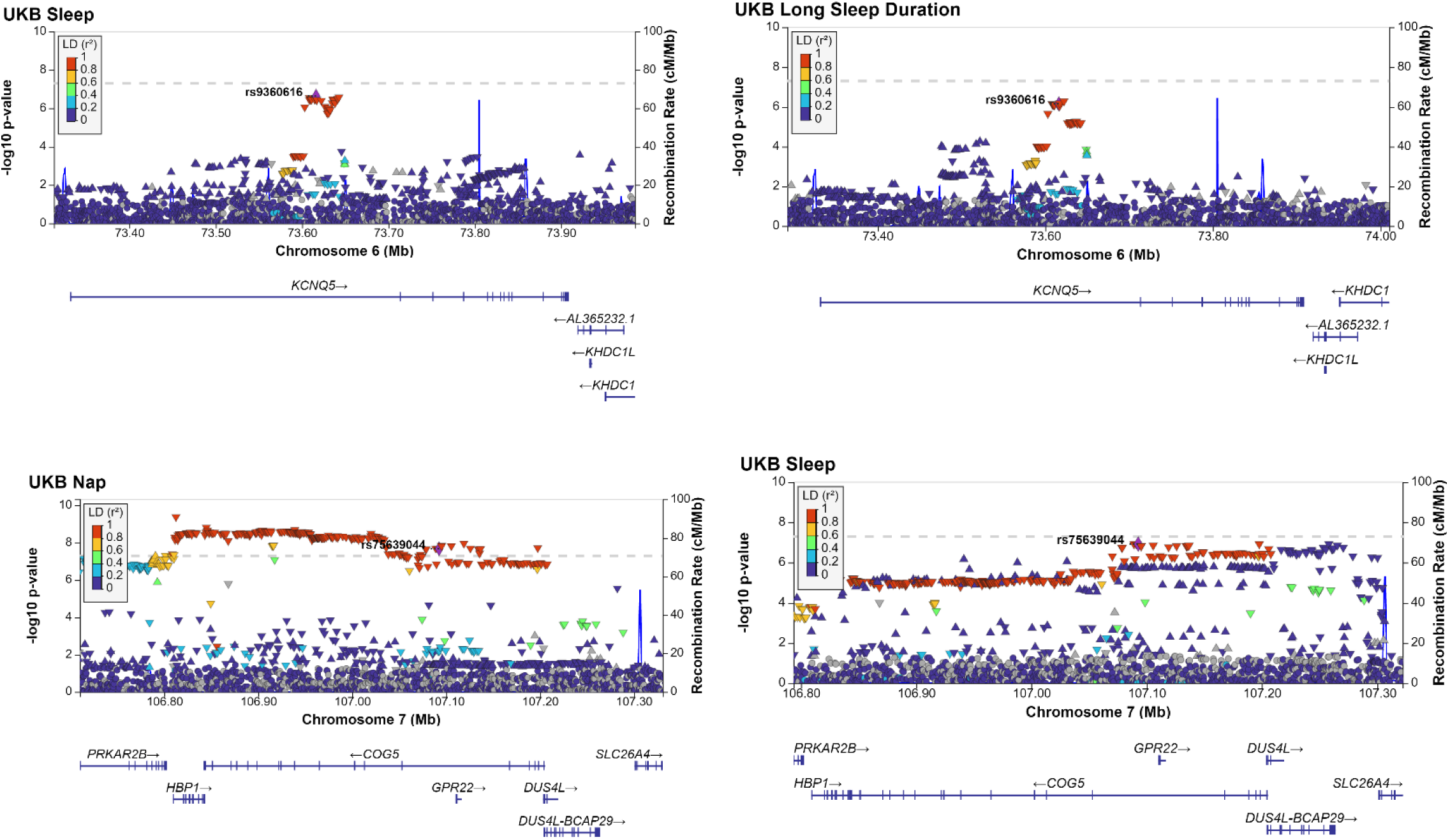
Two novel loci Colocalization with other sleep traits. Two novel loci in EUR ancestry GWAS (n=446,118). The novel loci (on chromosomes 6 and 7) are shown. Plots are centered on lead variants with 400 kb upstream and downstream from the lead variant. European ancestry LD was chosen as the LD reference for all plots (Methods).

